# LINES between species: Evolutionary dynamics of LINE-1 retrotransposons across the eukaryotic tree of life

**DOI:** 10.1101/050880

**Authors:** Atma M. Ivancevic, R. Daniel Kortschak, Terry Bertozzi, David L. Adelson

## Abstract

LINE-1 (L1) retrotransposons are dynamic elements. They have the potential to cause great genomic change due to their ability to ‘jump’ around the genome and amplify themselves, resulting in the duplication and rearrangement of regulatory DNA. Active L1, in particular, are often thought of as tightly constrained, homologous and ubiquitous elements with well-characterised domain organisation. For the past 30 years, model organisms have been used to define L1s as 6-8kb sequences containing a 5’-UTR, two open reading frames working harmoniously in *cis*, and a 3’-UTR with a polyA tail.

In this study, we demonstrate the remarkable and overlooked diversity of L1s via a comprehensive phylogenetic analysis of over 500 species from widely divergent branches of the tree of life. The rapid and recent growth of L1 elements in mammalian species is juxtaposed against their decline in plant species and complete extinction in most reptiles and insects. In fact, some of these previously unexplored mammalian species (e.g. snub-nosed monkey, minke whale) exhibit L1 retrotranspositional ‘hyperactivity’ far surpassing that of human or mouse. In contrast, non-mammalian L1s have become so varied that the current classification system seems to inadequately capture their structural characteristics. Our findings illustrate how both long-term inherited evolutionary patterns and random bursts of activity in individual species can significantly alter genomes, highlighting the importance of L1 dynamics in eukaryotes.

## Introduction

Transposable elements (TEs) are repetitive DNA sequences found in genomes scattered across the tree of life, and are often called ‘jumping genes’ because of their ability to replicate and move to new genomic locations. As such, they provide an important source of genome variation at both the species and individual level (Lynch 2006). Eukaryotic TEs are categorised based on their mechanism of retrotransposition. Class I retrotransposons use a copy-and-paste mechanism via an RNA intermediate, allowing massive amplification of copy number, which has the potential to cause substantial genomic change. Class II DNA transposons are more restricted because of their cut-and-paste mechanism. Retrotransposons are further divided into elements with (LTR) and without (non-LTR) long terminal repeats. Non-LTR elements comprise long interspersed elements (LINEs) and short interspersed elements (SINEs). LINEs are autonomous because they encode their own proteins for retrotransposition, whereas SINEs are non-autonomous and depend (in *trans*) on LINE-expressed proteins.

Long interspersed element 1 (LINE-1 or L1) is a well-known group of non-LTR retrotransposons found primarily in mammals (Kazazian 2000). Given their presence in both plant and animal species, L1s are very ancient elements; and it is assumed that they are ubiquitous across eukaryotes. More importantly, they are one of the most active autonomous elements in mammals, covering as much as 18% of the human genome (Furano 2000, Lander et al. 2001) and accountable for about 30% through amplification of processed pseudogenes and *Alu* SINEs (Esnault et al. 2000, Dewannieux et al. 2003, Graham and Boissinot 2006). This means that L1s are major drivers of evolution, capable of wreaking havoc on the genome through gene disruption (Kazazian 1998), alternative splicing (Kondo-Iida et al. 1999) and overexpression leading to cancer development and progression (Chen et al. 2005, Kaer and Speek 2013).

In the literature, active L1s are defined as 6-8kb elements containing a 5’-untranslated region (5’-UTR) with an internal promoter; two open reading frames (ORF1 and ORF2) separated by an intergenic region; and a 3’UTR containing a polyA tail (Furano 2000) (see Figure 1). ORF2 is around 3.8kb in length, translating to a 150kDa protein (ORF2p) which encodes an apurinic endonuclease (APE) and reverse transcriptase (RT) necessary for retrotransposition. ORF1 is much smaller (1kb nucleotide sequence; ORF1p is only 40kDa) and thought to have RNA-binding functionality (Furano 2000, Cost et al. 2002). This widely accepted structure has been used for over 30 years to identify putatively active elements in mammalian genomes (Scott et al. 1987). More recently, however, L1s with significant structural variations have been discovered – to the extent that the current terminology on what constitutes an L1 seems inadequate and limiting.

**Fig. 1.**
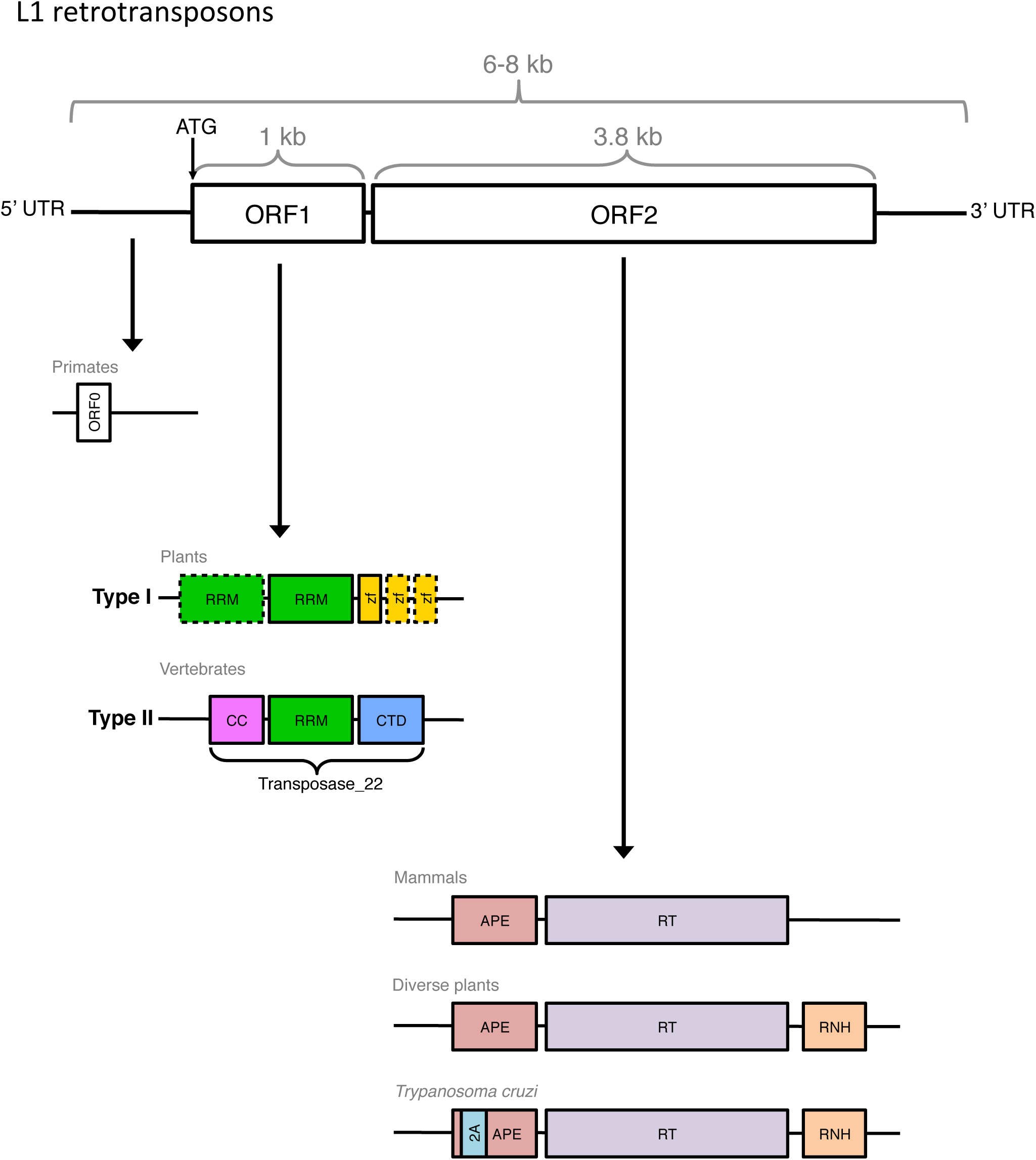
Conventional L1 structure and known variants. A functional L1 retrotransposon is 6-8kb in length and contains two ORFs, both of which encode proteins for retrotransposition. ORF0 has recently been discovered in primates and is thought to facilitate retrotransposition. L1 ORF1 sequences are divided into two types: Type II is widespread throughout vertebrates, while Type I has only been found in diverse plants. Likewise, domain variants of ORF2 have been found in some non-mammalian species (described in the main text). UTR, untranslated region; ORF, open reading frame; RRM, RNA recognition motif; zf, gag-like Cys_2_HisCys zinc knuckle; CC, coiled-coil; CTD, C-terminal domain; APE, apurinic endonuclease; RT, reverse transcriptase; RNH, ribonuclease H domain; 2A, oligopeptide translational recoding sequence.

For example, some plant species have been shown to contain an additional ribonuclease H domain (RNH) in ORF2p downstream of the RT domain, possibly acquired from domain shuffling between other L1s in plants, bacteria and Archaea (Smyshlyaev et al. 2013). The domains located within ORF1p can also vary drastically. Khazina and Weichenrieder (2009) classified retrotransposon ORF1ps into 5 types based on the presence and grouping of different domains, and indicated in which species/transposons each type was most commonly found. Type I ORF1p contains at least one RNA recognition motif (RRM) with a Cys_2_HisCys (CCHC) zinc knuckle, and is found in some plant L1s. Type II is the typical mammalian L1 ORF1p ‘Transposase 22’ (Finn et al. 2010), consisting of a coiled-coil (CC), single RRM and C-terminal domain (CTD). Type III and IV ORF1s are supposedly restricted to archaic elements such as CR1s (Chicken repeat 1) (Kapitonov and Jurka 2003) and L2s (Nakamura et al. 2012) and Type V are unclassified. However, even these classifications are insufficient. Metcalfe and Casane (2014) found that Jockey superfamily elements (especially CR1s and L2s) contain every possible type described by Khazina and Weichenrieder (2009), as well as further subtypes. This raises the question of whether L1s are also diverse in their structure, rather than being confined to Type II or I.

Some L1s do not appear to have an ORF1 region (Odon et al. 2013). For a long time, it was thought that co-expression of both ORF1p and ORF2p in *cis* was necessary for retrotransposition (Moran et al. 1996). However, L1 copies containing a disrupted ORF1p but intact ORF2p retain the ability to mobilise SINEs within the genome, as shown by Dewannieux et al. (2003) with a defective ORF1p mutant. Furthermore, some groups of non-LTR retrotransposons that only encode ORF2p are still capable of their own retrotransposition. Heras et al. (2006) discovered that *L1Tc* elements within the genome of *Trypanosoma cruzi* are able to retrotranspose without ORF1 due to the acquisition of a short 2A oligopeptide translational recoding sequence in the N-terminal region of ORF2 (Heras et al. 2009, Luke et al. 2013, Odon et al. 2013). Originally found only in RNA virus genomes, 2A sequences have now been shown to work in all eukaryotic cell types tested, mediating a translational recoding event known as ‘ribosome skipping’ (de Felipe et al. 2006).

Perhaps most intriguingly of all, recent evidence suggests the possibility of a third ORF in L1 elements: ORF0, an antisense open reading frame upstream of ORF1 (Denli et al. 2015). This ORF0 is very short, encoding a 71 amino acid peptide, and is thought to be primate-specific. Overexpression of ORF0p leads to a significant increase in L1 mobility, which may help explain the high retrotransposition activity of L1 in some primates (e.g. humans).

Growing evidence suggests that the current model of L1 activity (i.e. ORF1p + ORF2p in *cis* = retrotransposition) fails to capture the variation between different species or organisms, particularly beyond the mammalian lineage (Figure 1). In the past, detailed studies of the evolutionary dynamics of a transposable element were limited by the available genomes, which were mainly model organisms. This means that the evolutionary history of human and mouse L1 lineages have been extensively investigated, but there has not been much reported on most other mammalian groups, and even less for eukaryotes that are not mammals. In this study, we provide a definitive and comprehensive phylogenetic analysis of L1 content and activity in over 500 species from widely divergent branches of the tree of life. The genomes selected include plants, arthropods, reptiles, birds, mammals and other, more primitive eukaryotic species. We also include several cases of closely related organisms (within the same genus or species) to look for L1 differences between individuals, and the effects of different genome assembly methods. For each genome, we searched for the presence of L1 elements; and if found, characterised the elements as active or inactive and identified the domains in each of the ORF proteins. Our findings effectively illustrate the overlap between inherited evolutionary patterns and random individual bursts of activity, allowing a much broader understanding of transposable element dynamics in eukaryotes.

## Material and Methods

### Extraction of L1 Repeats from Taxa with Full Genome Data

Almost all of the genomes used in this study (499 out of 503) are publicly available from the National Center for Biotechnology Information (NCBI) (Sayers et al. 2012) or UCSC Genome Browser (Kent et al. 2002). Supplementary Table 1 lists the systematic name, common name, version, source and submitter of each genome assembly, and marks which genomes were privately acquired. If there was both a GenBank and RefSeq version for the genome, the GenBank version was used by default. Supplementary Table 2 shows the total genome sequence length and scaffold/contig N50 values, giving an approximation of the assembly quality. Since this study focused on analysis of full-length (~6kb), potentially active L1 elements, genome assemblies where scaffold number was an issue (e.g. causing hang-ups or taking too long) were filtered to remove scaffolds < 4kb. A phylogenetic representation of the genomic dataset was inferred using Archaeopteryx (Zmasek 2015) to download the Tree of Life (Maddison and Schulz 2007) topology for all Eukaryota (node identifier 3, ~ 76,000 species). The tree was extended (e.g. descendants added where necessary) to include all of the 503 genomes, and species not included in this study were removed. Out-dated branches were changed using OrthoDB (Kriventseva et al. 2015), OrthoMaM (Douzery et al. 2014), NCBI Taxonomy (Sayers et al. 2012) and recent publications (Murphy et al. 2001, Beck et al. 2006, Janecka et al. 2007) as references.

L1 hits were identified in each genome using an iterative query-driven method based on sequence similarity, as seen in Walsh et al. (2013). The original query sequences were obtained from Repbase (Jurka et al. 2005) or from past analyses (Adelson et al. 2009, Adelson et al. 2010). All of the accumulated query sequences were concatenated into one file, which was used as the input query to run LASTZ v1.02.00 (Harris 2007) with at least 80% length coverage. BEDTools v2.17.0 (Quinlan and Hall 2010) was used to merge overlapping hit intervals from different queries and extract a non-redundant set of L1 sequences in FASTA format. For each genome, the output hits were globally aligned with MUSCLE v3.8.31 (Edgar 2004) to produce a species consensus with Geneious v7.0.6 (Kearse et al. 2012). Genomes with a substantial number of hits required clustering with UCLUST v7.0.959_i86linux32 (Edgar 2010) before aligning. The species consensus sequences were then added to the query file (Supp. Figure 2).

This process was repeated three times, to accommodate inclusion of new genomes at various stages in the pipeline and to include diverse L1s to the set of queries. The last round of extraction used 643 L1 query sequences from 141 different species (including plants, animals, insects etc.). This query file is supplied in the Supplementary Material. Supplementary Table 5 shows the results from the final round of extraction, including the number of unique (merged) hits in each genome. Sample code for each step is available online (github.com/AdelaideBioinfo).

### Identification of Putatively Active L1 Elements

BEDTools was used to extend each L1 hit by 1kb either side before the ORF analysis, to overcome incomplete 5’ and 3’ ends that may be missing crucial start/stop codons. Geneious was then used to scan for potentially active L1s that met the length requirements. An L1 was defined as an ‘active’ candidate if it contained at least an intact ORF2 (regardless of the state of ORF1), as this means that it is either fully capable of retrotransposing itself (Moran et al. 1996, Heras et al. 2006) or it can cause activity in the genome by mobilising SINEs (Dewannieux et al. 2003). An ORF was considered intact if it was at least 80% of the expected length (≥ 800bp for ORF1 and ≥ 3kb for ORF2 – Supp. Figure 4). ORF1 had the additional requirement of having a methionine (ATG) as the start codon (Penzkofer et al. 2005). ORF1s were further confirmed by manually checking their position in each species global alignment (this was necessary because in a large number of species, the ORF2 was truncated to the extent that it only contained an endonuclease region of about 1kb, which could be mistaken for the ORF1). Finally, all putative ORF1 and ORF2 amino acid sequences were checked for similarity to known domains using HMM-HMM comparison (Finn et al. 2011) against the Pfam 28.0 database (Finn et al. 2010) as at May 2015 (includes 16230 families).

### Dendrogram Construction from Nucleotide L1 Sequences

Full-length L1 hits (or near full-length, for species with low copy number) were globally aligned using MUSCLE (Edgar 2004). Mammalian species required clustering with UCLUST (Edgar 2010) before aligning, due to the huge number of hits. Supplementary Table 4 shows the clustering percent identity used for each species to produce alignable clusters/families. Sequences in the alignment were tagged (e.g. a label was appended to the sequence names) to indicate which ORFs were considered to be intact. The dominant active clusters for each species were represented as dendrograms, or unrooted tree diagrams, using FastTree v2.1.7 (Price et al. 2010). Archaeopteryx v0.9901 beta (Zmasek 2015) was used to visualise and annotate each tree based on their sequence name tags. Dendrograms constructed for each species are included in the Supplementary (PDF and Newick tree format).

### Phylogenetic Analyses of Translated Open Reading Frames using a Single Consensus Approach

All confirmed ORF2 sequences in each species were extracted, translated into amino acid residues and globally aligned with MUSCLE (Edgar 2004) (Supp. Figure 5). The consensus for each species was generated in Geneious (Kearse et al. 2012) using majority rule (most common bases, fewest ambiguities) and a base was regarded ambiguous if coverage at that position was < 3 sequences (unless the alignment had ≤ 3 sequences, in which case this was changed to < 2 sequences. Baiji (Chinese dolphin), mouse and elephant were also changed to < 4 sequences, due to the large number of sequences). This produced a single L1 ORF2p consensus for each species. These consensus sequences were globally aligned using MUSCLE (Edgar 2004) and a phylogeny was inferred with maximum likelihood using FastTree (Price et al. 2010), which uses a general time reversible (GTR) model and gamma approximation on substitution rates. The same process was repeated to generate an ORF1p species consensus tree (provided as Supp. Figure 9 because it is not as accurate as the ORF2p tree).

### Clustering Analysis of L1 Proteins ORF1p and ORF2p

All HMMer-confirmed ORF1p and ORF2p amino acid sequences were extracted and clustered (separately for each ORF type) using an all-against-all BLAST (Altschul et al. 1990) approach. The BLAST was performed using BLAST v2.2.24 and NCBI-BLAST v2.2.27+ (Altschul et al. 1990) with the following parameters:-p blastp,-e 1e-10,-m 8 (for tabular output). Based on the BLAST results, the ORFs were then clustered using SiLiX software (Miele et al. 2011) with default parameters and–net to create a net file which contains all the pairs taken into account after filtering.

## Results

### Ubiquity of L1 Across the Eukaryotic Tree of Life

To simplify discussion of the results, we define three different states that a genome can be in in terms of L1 content: absent (L1^−^), meaning that no L1s were detected in the genome; present (L1^+^), meaning that L1s were found in partial or full-length form; and active (L1*), meaning that at least one putatively active L1 was found in the genome. L1^−^ and L1^+^ are mutually exclusive (a genome cannot have both presence and absence of L1s), while L1* is the active subset of L1^+^. Using this ternary system, we screened 503 eukaryotic species representing key clades of the tree of life (123 plants, 145 protostomes, 98 mammals, 74 reptiles and birds, 22 neopterygians, 11 flatworms and 30 other species) (Figure 2, Supp. Figure 1 includes tip labels). Of these, 335 genomes were found to be L1^+^. L1 copy number was highest in mammals, with hundreds to thousands of full-length L1 sequences found in almost every mammalian species analysed (with the exception of monotremes: platypus only contained a small number of fragmented L1s, and echidna did not reveal any signs of L1s, possibly due to the draft quality of the assembly).

**Fig. 2.**
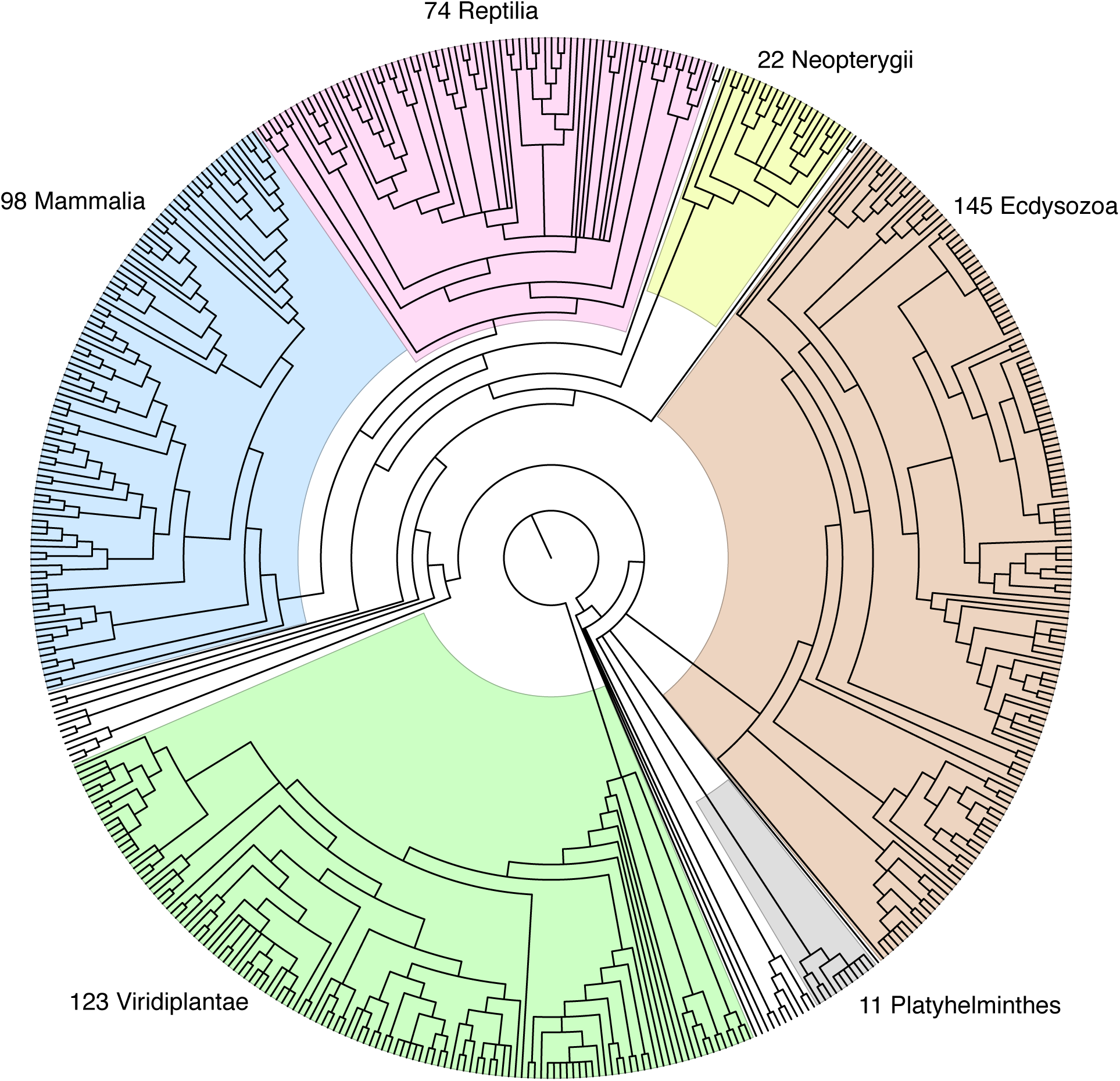
Phylogenetic representation of genomic dataset. Species relationships between the 503 representative genomes used in this study were inferred using Archaeopteryx to download the Tree of Life topology for all Eukaryota (node id 3) and extract the 503 species of interest. Out-dated branches were updated using OrthoDB, OrthoMaM, NCBI Taxonomy and recent publications as references. Labels indicate the major groups present in this dataset.

L1s also appeared frequently in plants (107/123 L1^+^ plant species), but colonised far less of each genome (e.g. typical copy number of 10-100 L1s). Fish, reptiles and amphibians had similarly low copy numbers and a patchy distribution. Birds had an exceptionally low (yet consistent) L1 copy number: only 1 L1 element was found in most of the bird species analysed, yet this element was conserved through enough species that it is likely an ancient remnant of L1 from a common ancestor.

In the protostomes, L1s were almost completely absent except in mosquitos: all three of the studied mosquito species contained full-length hits. Some other, more primitive genomes had a surprising number of L1s: three different species of sea squirts and a sea anemone all had full-length or near full-length hits. Supplementary Table 5 contains a summary of the L1 hits found in each genome and the size range of the hits.

### Dead Or Alive - How Many L1s Have Retained Their Activity?

Of the 335 L1^+^ eukaryotes, 125 species were further determined to be L1*: 62 plants, 53 mammals, and 10 non-mammalian animal species. This is illustrated in Figure 3 (mammals) and Figure 4 (plants), which highlight the species where L1s are present or active (full tree available as Supp. Figure 6). Although all coloured branches indicate presence (L1^+^), the active subset (L1*) is coloured magenta, so in this case the blue branches (L1^+^ – L1*) indicate species that only contain ‘extinct’ L1s (i.e. present but inactive). Since the L1 state of each genome is only observable at the tree tips, the phylogeny was annotated based on the most parsimonious explanation being that a loss of activity is more likely than a gain (hence ancestral branches are coloured ‘active’ if any of the descendants display activity). Noticeably, despite the ubiquitous presence of L1 across the mammalian lineage, L1 in quite a few mammalian species or subgroups (e.g. megabats, some rodents and Afrotherian mammals) are no longer active. In contrast, other mammals seem to be bursting with L1 activity: including several species (e.g. minke whale, antelope, snub-nosed monkey, panda, baiji) which have not been studied before in the context of L1 retrotransposition. Snub-nosed monkey (*Rhinopithecus roxellana*) is a particularly good example because human is often thought of as the most retrotranspositionally-active primate - even though only 6 ‘hot’ L1s contribute to the majority of retrotransposition in the human genome (Brouha et al. 2003). Snub-nosed monkey, on the other hand, contains 2887 putatively active L1 sequences, suggesting that its retrotransposition potential is substantially higher than that of human or any other primate.

**Fig. 3.**
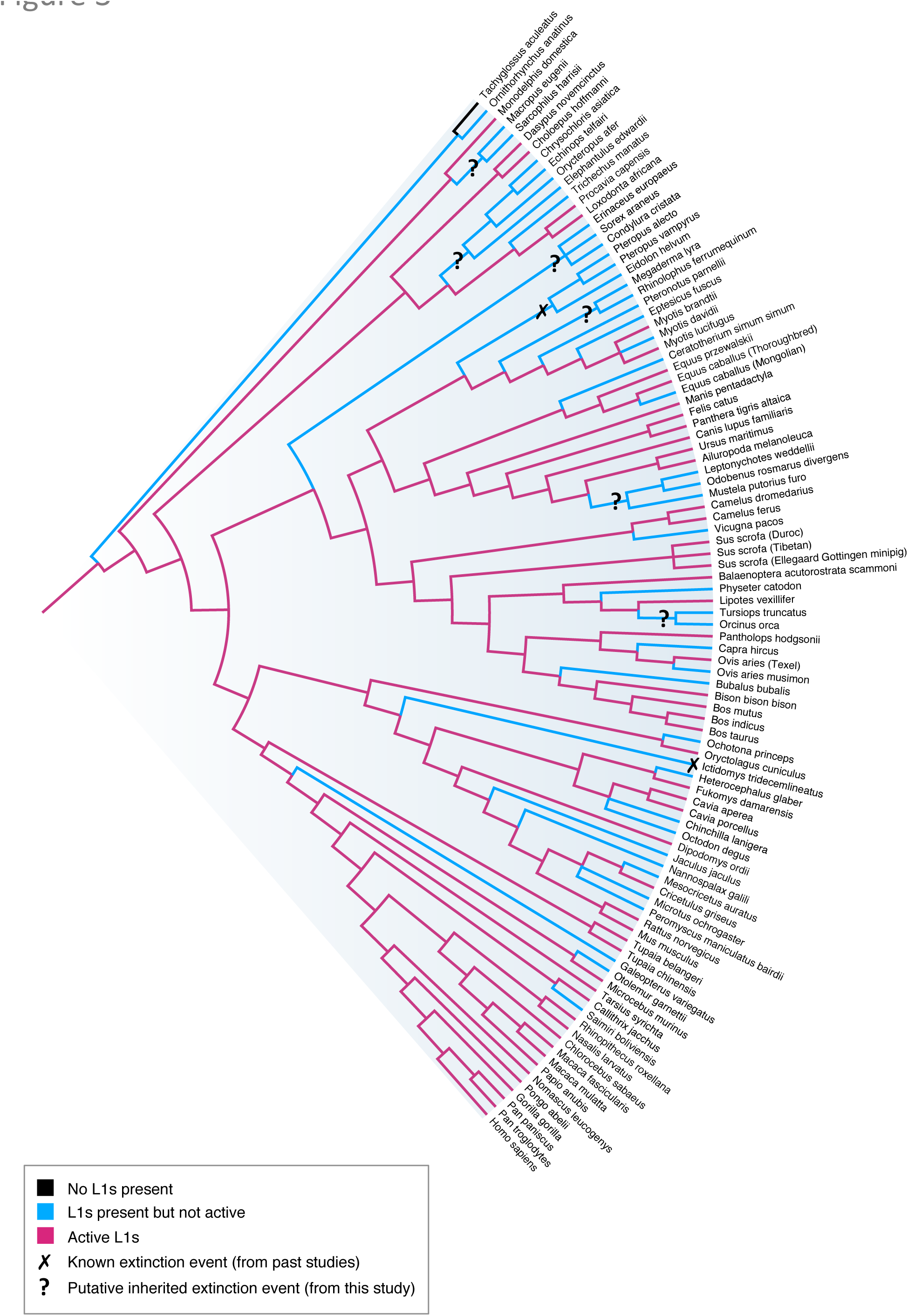
Mammalian phylogeny reveals ubiquitous L1 presence and multiple possible extinction events. Mammalian subset of the inferred tree of life showing how genomes are classified as L1 absent (L1^−^) (black), L1 present but inactive (L1^+^ – L1*) (blue) or L1 active (L1*) (red). Putative extinction events that have not been previously investigated (marked by ‘**?**’) are only included if at least two monophyletic species exhibit L1 extinction.

**Fig. 4.**
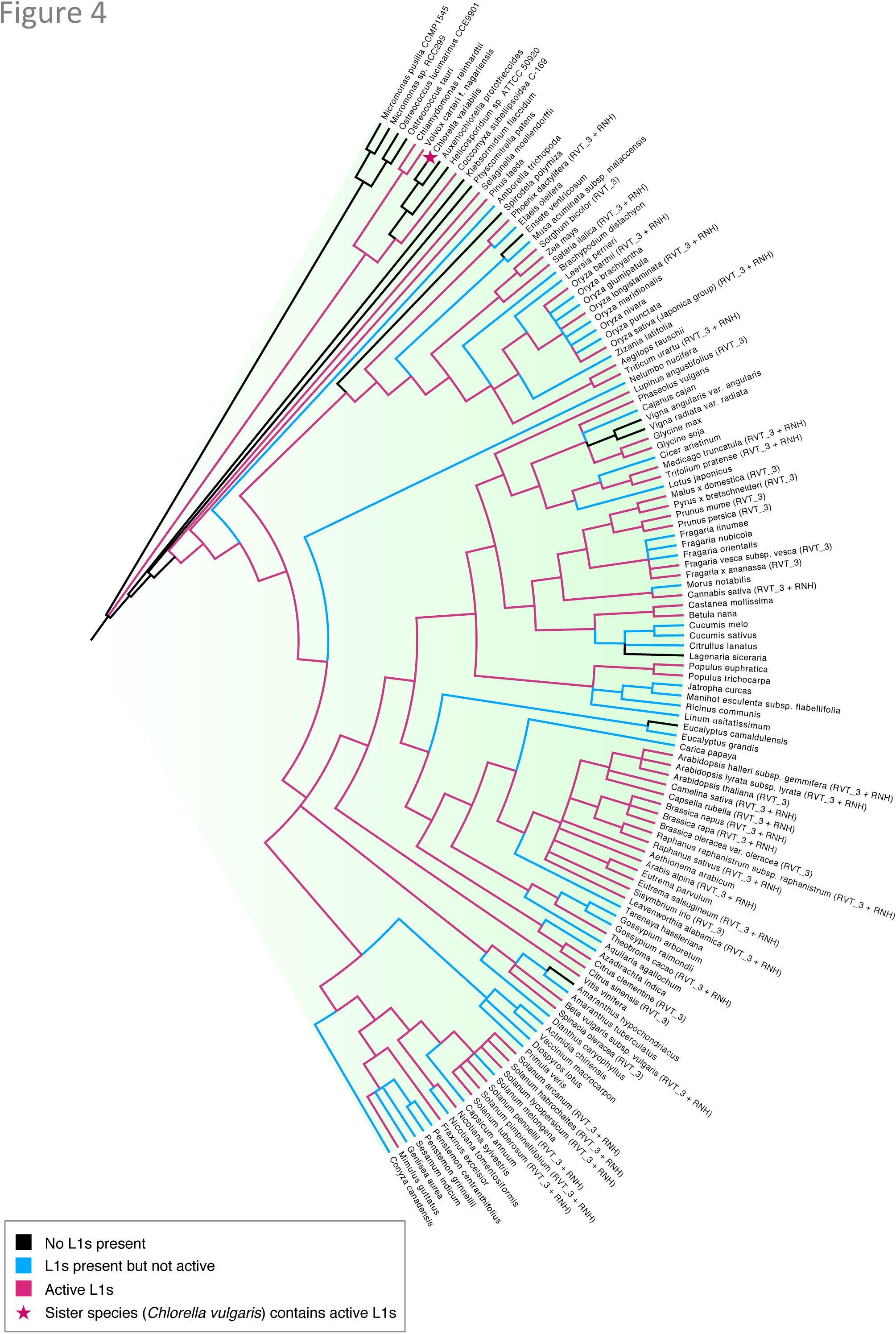
Plant phylogeny showing the sporadic distribution of active L1s. Plant subset of the tree of life showing the L1 state of each genome (coloured branches), and whether or not the genome contains a plant-specific reverse transcriptase (RVT_3) or ribonuclease H domain (RNH) (annotations appended to the tip labels). Brassicales stands out as the most dominant L1* family, and almost all of the species in the family contain RVT_3 + RNH ORF2p domains. This tree also highlights an apparent discrepancy: *Coccomyxa subellipsoidea*, which contains a novel HTH_1 ORF1p domain and exhibits 100% retrotranspositional potential, is placed next to an L1-lacking subgroup of species.

Beyond the mammalian lineage, there seem to be sporadic bursts of L1 activity. For example, there are only two L1* fish species, one frog, one lizard and a few mosquitos. The Atlantic canary genome (*Serinus canaria*) appeared to be the only L1* bird species; but upon closer inspection, it was found that this L1 hit came from an isolated 4kb scaffold that contained a human-like (over 98% similarity) ORF2. Furthermore, over 1000 other short scaffolds (<500bp in length) in this genome were classified by Repbase (Jurka et al. 2005) as belonging to human, so the L1 was disregarded as likely contamination.

In plants, the L1 state of species looks much more sporadic. *Brassicales* stands out as the most dominant family, with each member bearing a significant number of active L1s. Another notable L1* species was *Coccomyxa subellipsoidea*, which only contained 15 L1 elements but every single one of these elements is putatively active and almost identical, suggesting recent retrotransposition. This genome also appears as a discrepancy in our tree; it is the only instance where a L1* species is phylogenetically placed next to a L1^−^ species (see Figure 4). However, given that our dataset does not contain all species, this could be a result of incomplete sampling and hence incorrect placement of the species. Accordingly, the ancestral branch was coloured red (L1*) despite the absence of L1s in several descendent species, because another study shows that *Chlorella vulgaris* (sister to *Chlorella variabilis)* contains active L1-like elements 98% identical to *Coccomyxa subellipsoidea* (Higashiyama et al. 1997).

Finally, the number of active L1s found in each genome was compared to the total number of full-length L1s in that genome, to get a percentage estimate of L1 activity per species (Figure 5, Supplementary Table 7). We found that mammalian species often contain a large number of inactive elements, so the percentage of active L1s is relatively low (e.g. < 20%). A similar pattern is observed in plants, albeit at lower copy number. However, in the 10 non-mammalian animal species, a significant number of L1s in the genome appear to be active L1s (despite the low copy number). Consequently, the centroid of the graph is shifted to the right, with all of the non-mammalian species (e.g. mosquitos, fish, frog, lizard, sea squirt) having more than 19% of their L1s putatively active.

**Fig. 5.**
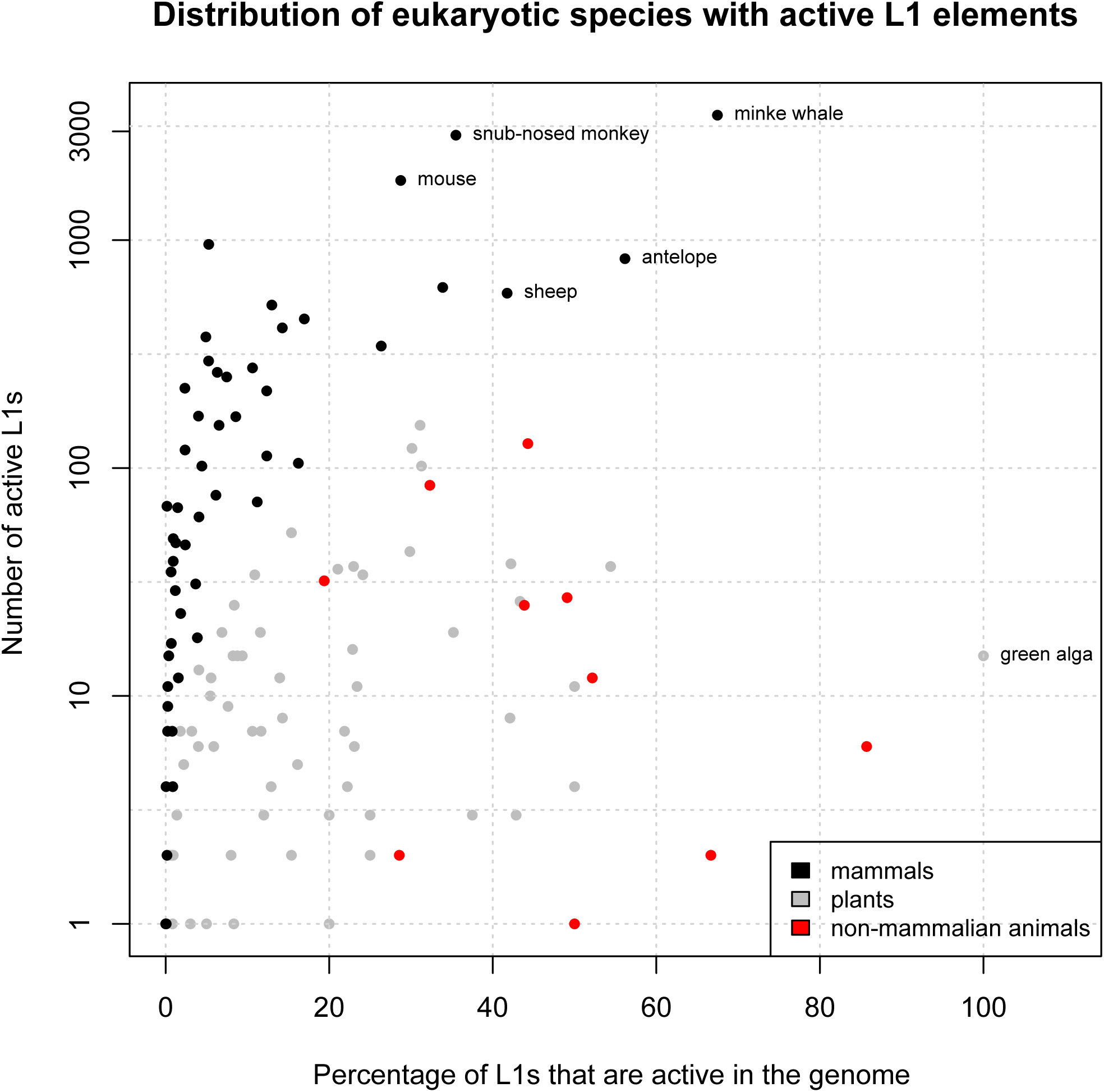
Distribution of active L1 elements reveals several ‘hyperactive’ mammalian species. Non-mammalian animal species (red) and plants (grey) appear to have high retrotranspositional potential but low observable L1 activity in the genome (low copy number). In contrast, mammals (black) typically have a very high L1 copy number, but the majority of these are inactive. The labelled mammalian species stand out as L1 ‘hyperactive’ species because they are the most likely to be currently replicating and expanding within the genome.

### Mammalian Species Typically Have A Dominant Active Cluster

The clustering and dendrogram construction of L1 nucleotide sequences revealed that most mammals contain one large, dominant active cluster of closely related elements. As mentioned above, snub-nosed monkey is a remarkably active species in a comparatively inactive subgroup (i.e. primates). It is also the only species where the main cluster of L1s was so large that it had to be clustered at 90% identity to be alignable (all other mammalian species were clustered at 70, 80 or 85% - see Supp. Table 4). This resulted in the final cluster containing 3303 full-length L1s with 94.8% pairwise identity (Supp. Table 8), which was used to construct an unrooted maximum likelihood tree (Figure 6a,b) highlighting elements with ORF1 intact, ORF2 intact, or both ORFs intact. Almost all of the L1s in this cluster have both ORFs intact (Figure 6b) and are clustered on the shorter branches, indicating very recent activity.

**Fig. 6.**
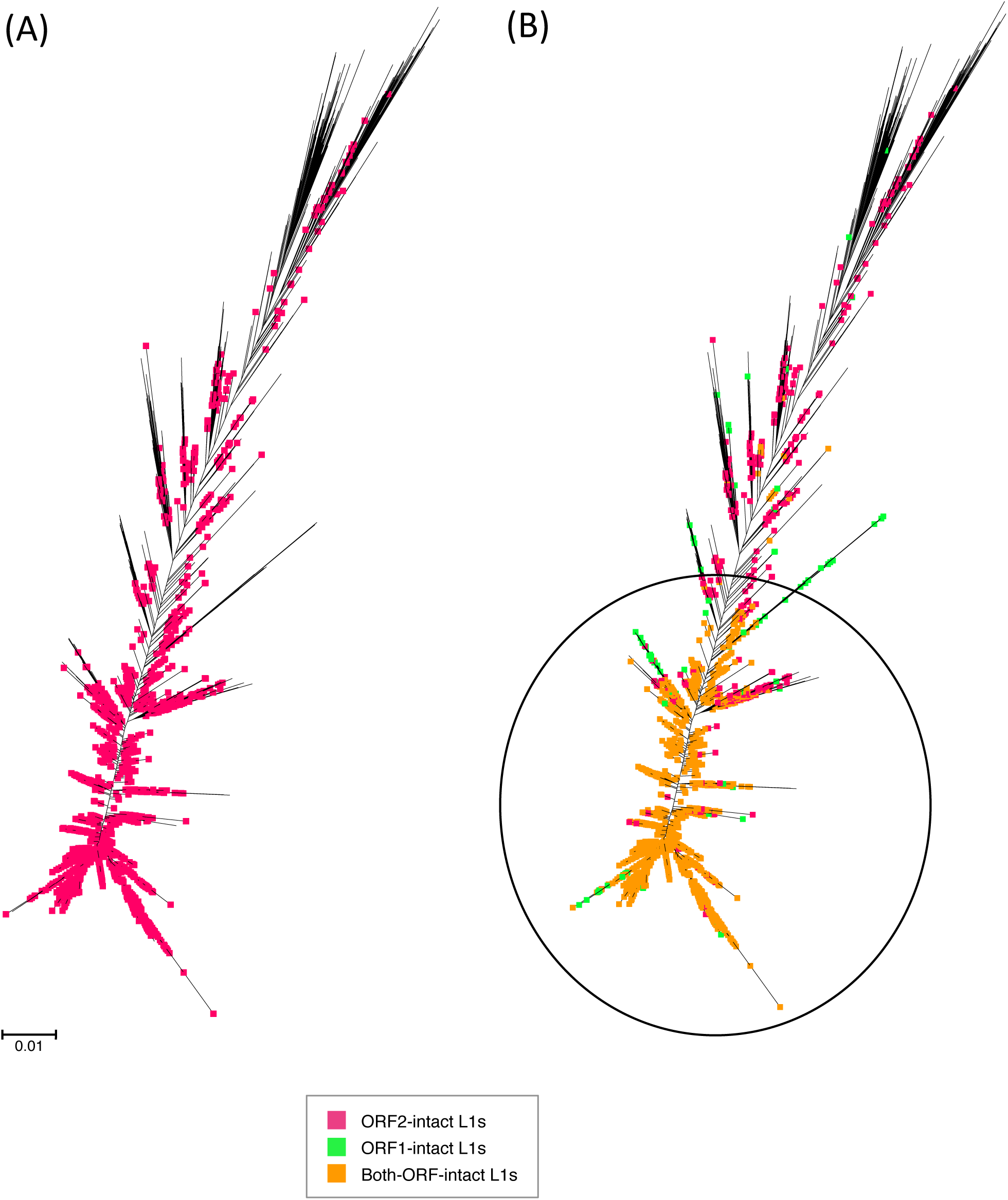
Master lineage model predominant in most mammalian species, including primate *Rhinopithecus roxellana*. Maximum likelihood dendrogram inferred using FastTree from full-length L1 nucleotide sequences extracted from full genome species data. Sequences were clustered with UCLUST and globally aligned with MUSCLE. Species with a clearly dominant L1* cluster were classified as master lineage models, as shown. Sequences in the alignment were tagged to indicate which ORFs were intact and visualised using Archaeopteryx.

However, in some species it is obvious that there is more than one significant active cluster. Horse (*Equus caballus*) is a well-known example of a species with five L1 (equine) subfamilies, two of which contain active elements (Adelson et al. 2010). Megabats are also known to have harboured multiple contemporaneous L1 lineages, although those lineages are now extinct (Yang et al. 2014). Nonetheless, this multiple lineage phenomenon seems to extend to the microbat subgroup as well: Figure 7 depicts the clustering and dendogram construction for *Myotis brandtii*, where there is no discernible dominant cluster. The elements in each cluster are >70% similar to each other, but the clusters themselves are distinct at this level. Once again, we see a tendency for active L1s to converge on the shorter branches; in fact, they are almost always restricted to these shorter branches, unlike in snub-nosed monkey.

**Fig. 7.**
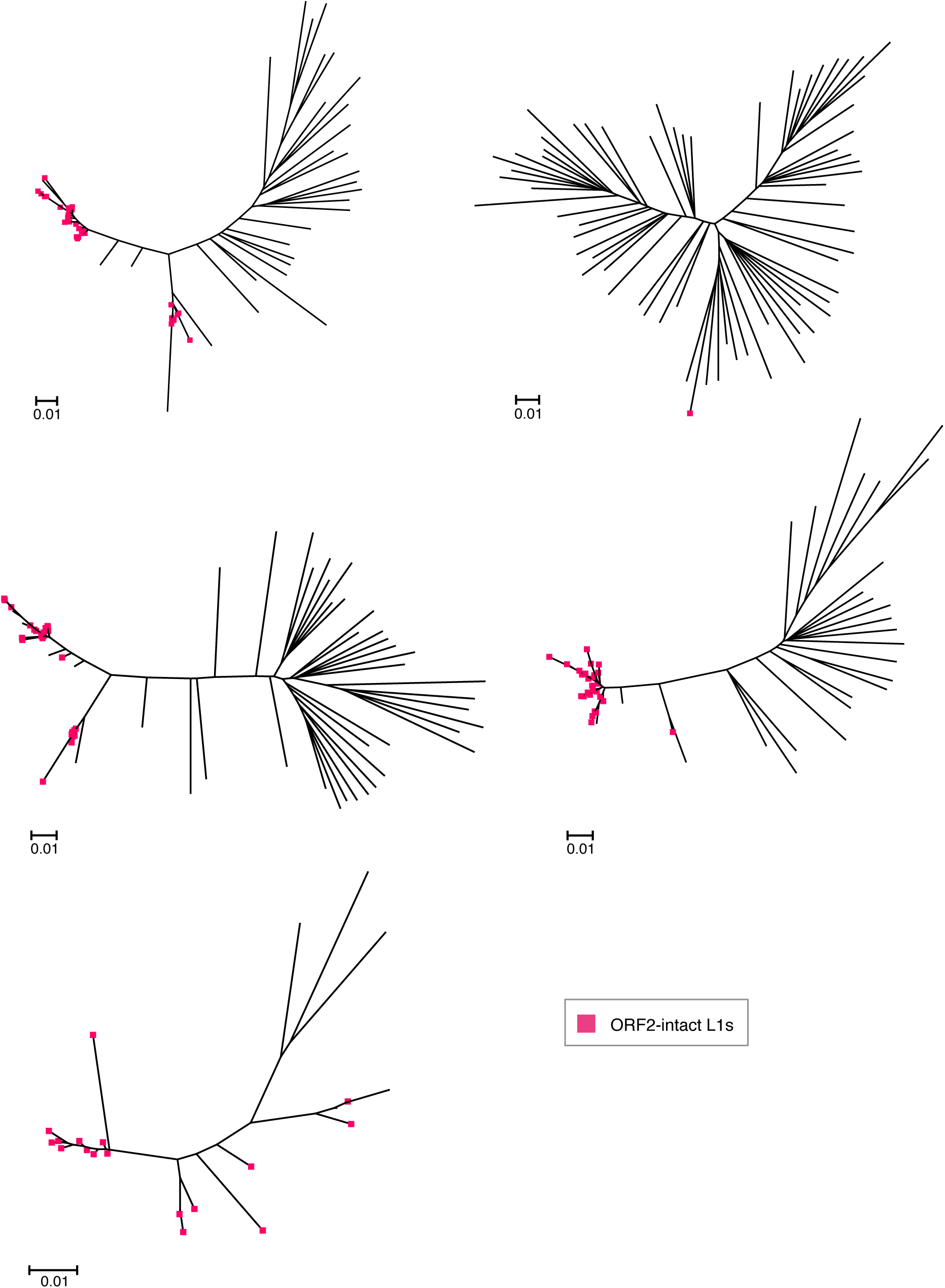
Multiple L1 lineages present in the *Myotis brandtii* genome. Maximum likelihood dendrogram inferred using FastTree from full-length L1 nucleotide sequences extracted from full genome species data. As in Fig. 6., sequences were clustered with UCLUST, aligned with MUSCLE, annotated with Geneious and visualised with Archaeopteryx. In this case, only ORF2-intact sequences are highlighted.

Even more unexpected are the species (e.g. opossum, sheep, cow, rabbit and some primates – see Supp. Table 8) where the clustering classifies the potentially active elements as ‘singletons’ or ‘unclustered’ sequences (Edgar 2010). This means that the active elements are less than 70% similar to each other: a scenario which suggests that (a) these elements are not really active, because they never expanded within the genome; (b) the L1 lineages are old enough to have accumulated substantial mutations; or (c) they have only just appeared in the genome, either through horizontal acquisition or a reversion mutation.

### Variation of ORF1p and ORF2p across Species

As expected, both ORF1p and ORF2p revealed significant variation between species. The main difference in ORF2p structure was the presence of a different reverse transcriptase (RVT_3 versus the expected RVT_1) (Finn et al. 2010) in 42 plant species; 28 of which also had an extra RNH domain (recently investigated by Smyshlyaev et al. (2013)). Interestingly, all of the most L1-active plant orders (e.g. Rosales, Brassicales, Solanales) seem to contain RVT_3, or RVT_3 + RNH (Figure 4). It is possible that most plant species have lost the ability to retrotranspose (due to an inability to utilise the normal RVT_1), but some species have acquired a plant-specific reverse transcriptase domain(s) and thus are still able to retrotranspose.

This extra RNH domain (along with multiple random insertions of other protein coding sequences), means that plant ORF2s often extended beyond the expected 3-3.8kb. Figure 8 (Supp. Figure 10 includes node support values) provides striking graphical evidence for this: plant species cluster on long branches and over multiple clades, indicating either a high mutation rate (supported by the domain analysis) or long periods of quiescence (also plausible, given the sporadic active/non-active patterns in Figure 4). In contrast, ORF2-intact animal species (particularly mammals) form one large all-inclusive clade with relatively short terminal branches, indicating recent and highly similar families. Looking at the pairwise identities between L1s in each species (Supp. Table 10) further reinforces the divergence of plant ORF2ps (30-50% pairwise identity) versus mammals (over 90% for most species). Nonetheless, the gross ORF2p tree topology reflects the expected species relationships; implying that there is selective pressure for the ORF2 structure to remain conserved enough to retain activity.

**Fig. 8.**
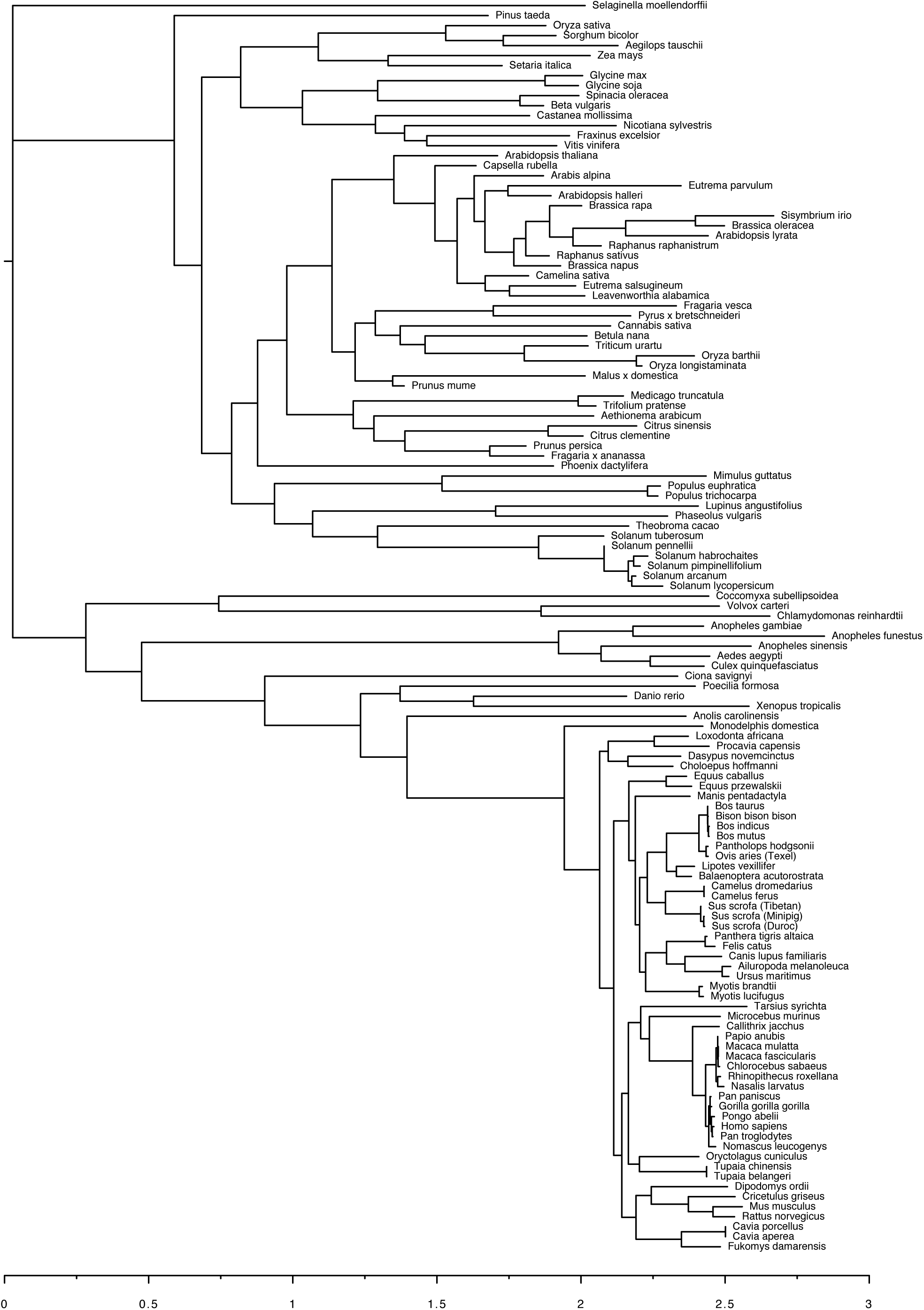
ORF2p consensus tree highlights differences between plant and animal L1s. Maximum likelihood phylogeny inferred using FastTree from L1 ORF2p consensus sequences extracted from genome data. ORF2 nucleotide sequences were identified and extracted with Geneious, translated to amino acid residues (ORF2p) and aligned with MUSCLE. The scale axis at the bottom of the figure accentuates the difference between animal ORF2p (closely related, recent retrotransposition) compared to plant ORF2p (diverse and comparatively ancient). Phylogenetic relationships between species appear conserved.

The variability found in ORF1 sequences, both within and between species, is even more staggering. Almost all of the ORF1-intact animal species (all 86 mammals and 10 non-mammalian animals) had Transposase_22 as the top hit for ORF1p (Figure 9a, Supp. Table 11), which corresponds to the Type II ORF1p defined by Khazina and Weichenrieder (2009) (Figure 1). Plants, mosquitos and seq squirt (*Ciona savignyi*), on the other hand, contain a domain structure that very loosely resembles that of Type I ORF1ps (Khazina and Weichenrieder 2009); but only because of the presence of RRM or zf-CCHC domains (Figure 9b). The most dominant plant domain was, in fact, an uncharacterised domain of unknown function (DUF4283) (Finn et al. 2010). This domain was present in 75 of the 78 ORF1-intact plant species, and in some species (e.g. *Vitis vinifera*) it was the only ORF1 domain present. Likewise, *Coccomyxa subellipsoidea* did not contain any domains previously associated with ORF1 proteins – instead, the entire ORF1 amino acid sequence for each of its L1s was classified as HTH_1 (Figure 9a): a bacterial regulatory helix-turn-helix protein of the LysR family (Finn et al. 2010). Further examination of this novel domain revealed that *Coccomyxa subellipsoidea* ORF1ps, as well as full-length L1 nucleotide sequences, are 98% identical to Zepp; a LINE-like retrotransposon found in *Chlorella vulgaris* (Higashiyama et al. 1997). *Chlorella vulgaris* was not included in this study as the assembly is only available in contig form. However, another *Chlorella* species (*C. variabilis*) was included and showed no L1 presence at all (Figure 4). Given that *Coccomyxa subellipsoidea* contains a typical ORF2p structure, it is possible that the abnormal ORF1p (or entire L1) was acquired from another species (e.g. *Chlorella vulgaris*). Alternatively, TEs have a tendency to take necessary proteins directly from their host (Abrusan et al. 2013), so the L1s may have utilised host-HTH_1 proteins to instigate retrotransposition.

**Fig. 9.**
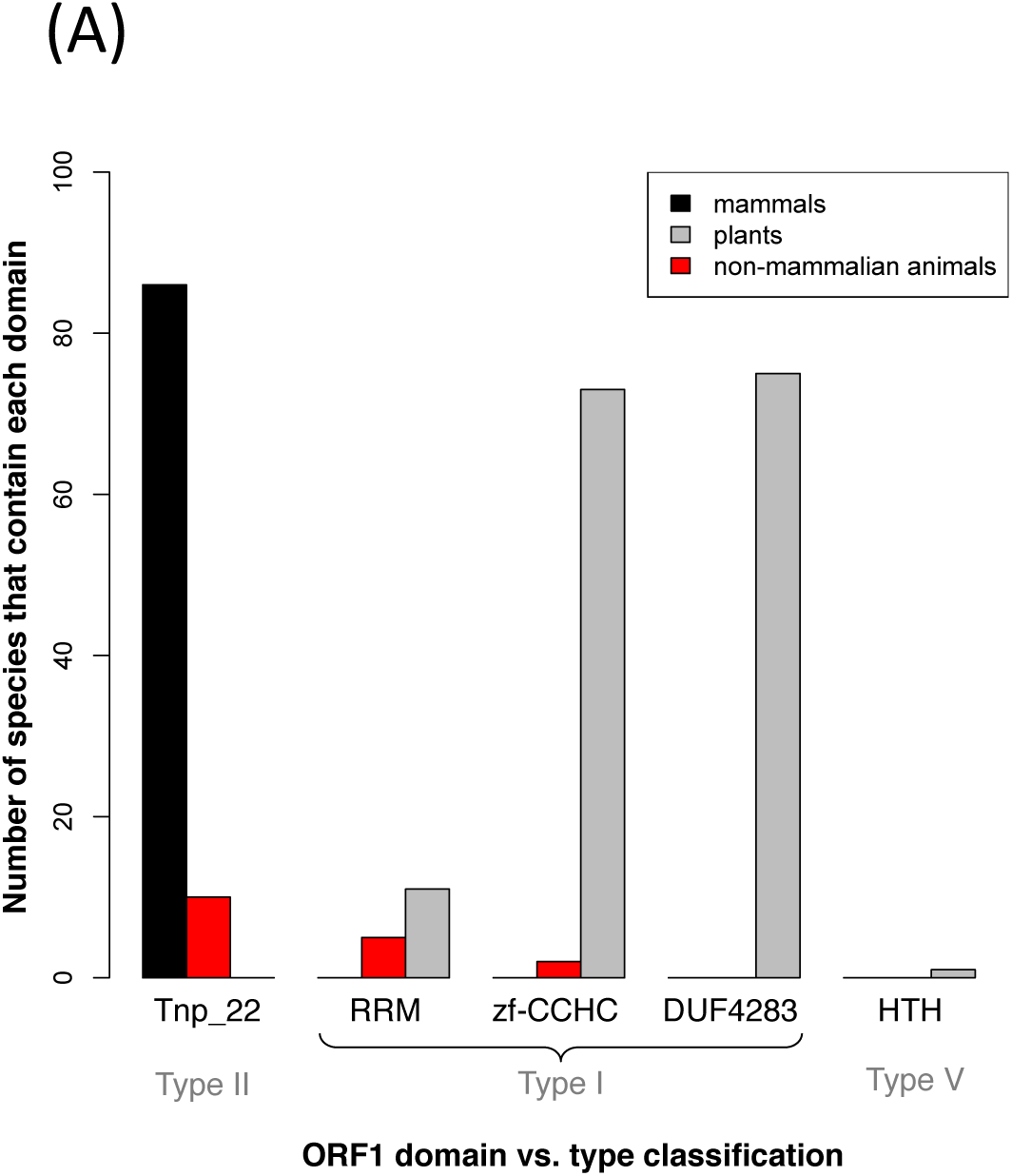
ORF1p clustering and domain identification analysis. **9a) ORF1p domain summary from HMM-HMM comparison**. Type II ORF1p contains a distinctive Transposase_22 domain. This domain was found in all 86 ORF1-intact mammalian species and 10 non-mammalian animal species. Type I ORF1p is indicated by presence of RRM/zf-CCHC domains. Based on this criteria, most of the plants (77/78) and 6 non-mammalian animals (mosquitos and sea squirt) contain Type I ORF1p. However, an uncharacterised plant domain (DUF4283) is the most prevalent ORF1p domain in plants. One plant species (*Coccomyxa subellipsoidea*) belongs to Type V (unclassified) because it contains a novel HTH_1 domain. Note that there are only 78 ORF1-intact plant species, yet most of them appear in multiple domain categories because they contain various domain combinations (e.g. DUF4283 + RRM, or DUF4283 + zf).

**9b).**
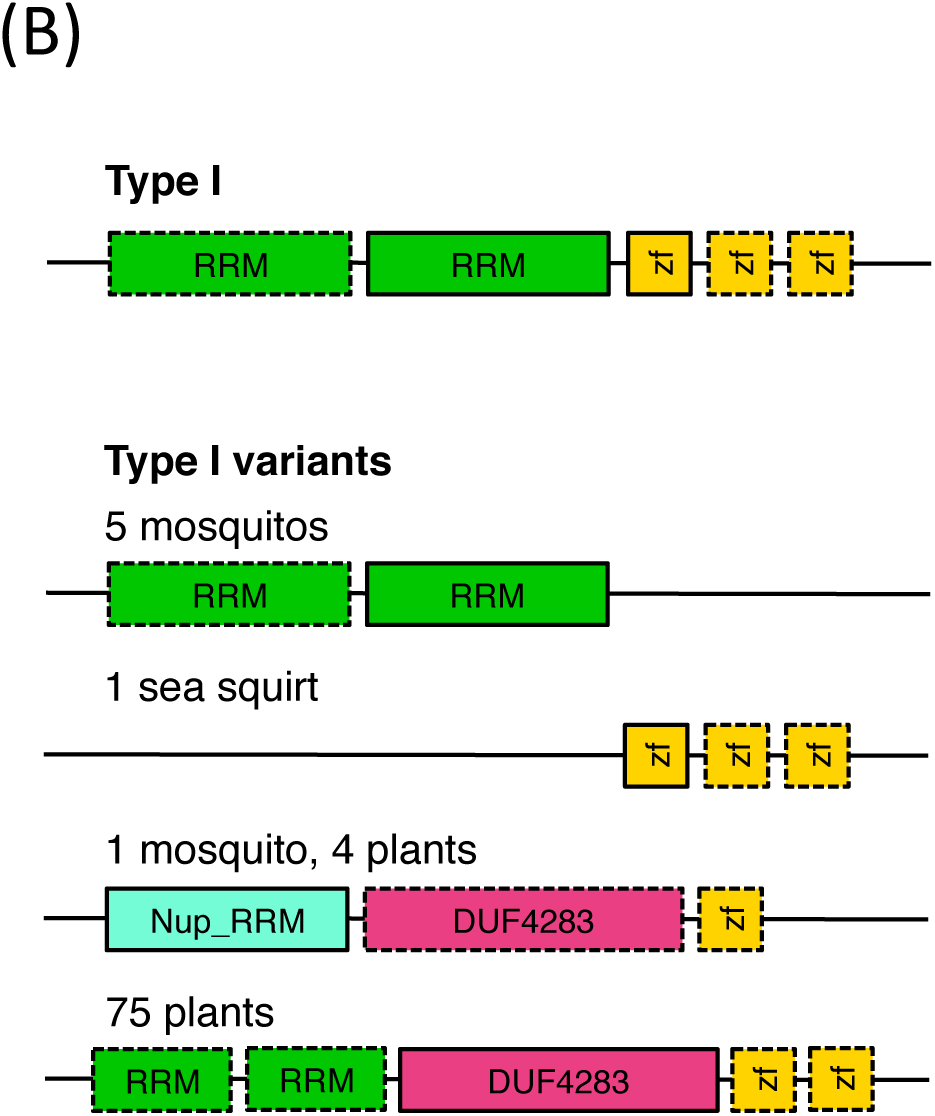
Variants of Type I ORF1 proteins. Type I ORF1p are characterised by the presence of at least one RRM and zf-CCHC domain. Significant variations to this type were found in plant and non-mammalian animal species, e.g. mosquitos lack zf-CCHC domain(s); sea squirt lacks RRM domain(s); several species contain an unusual RRM domain (Nup_RRM); and plants predominantly have an unknown DUF4283 domain, regardless of the RRM and zf-CCHC domains.

### Antisense Characteristics of Active L1s

The analysis of ORF1 and ORF2 sequences across genomes led to the discovery of an antisense open reading frame overlapping ORF1. This novel ORF was initially noticed in the panda genome (*Ailuropoda melanoleuca*), where it is present in almost every L1 element that has both ORFs intact (586/594). As a result, we screened each genome for strictly active L1s (i.e. both ORF1 and ORF2 intact) to determine whether other species contained similar antisense ORFs (i.e. overlapping ORF1 in the reverse direction and about 1kb in length). Apart from panda, only eight other mammalian species contained anything remotely similar (Figure 10a), albeit at lower copy number (Supp. Table 14). No such reverse ORFS were found in any of the non-mammalian animal or plant species. Interestingly, these ORFs only appeared in mammalian species with a substantial number of active L1s (e.g. minke whale, baiji, dog, rat), suggesting that they might somehow contribute to L1 retrotransposition; yet they are noticeably absent from all of the primates, including snub-nosed monkey. They are also clearly distinct from the primate-specific antisense ORF0 (Denli et al. 2015), which is much shorter and upstream of ORF1.

**Fig. 10.**
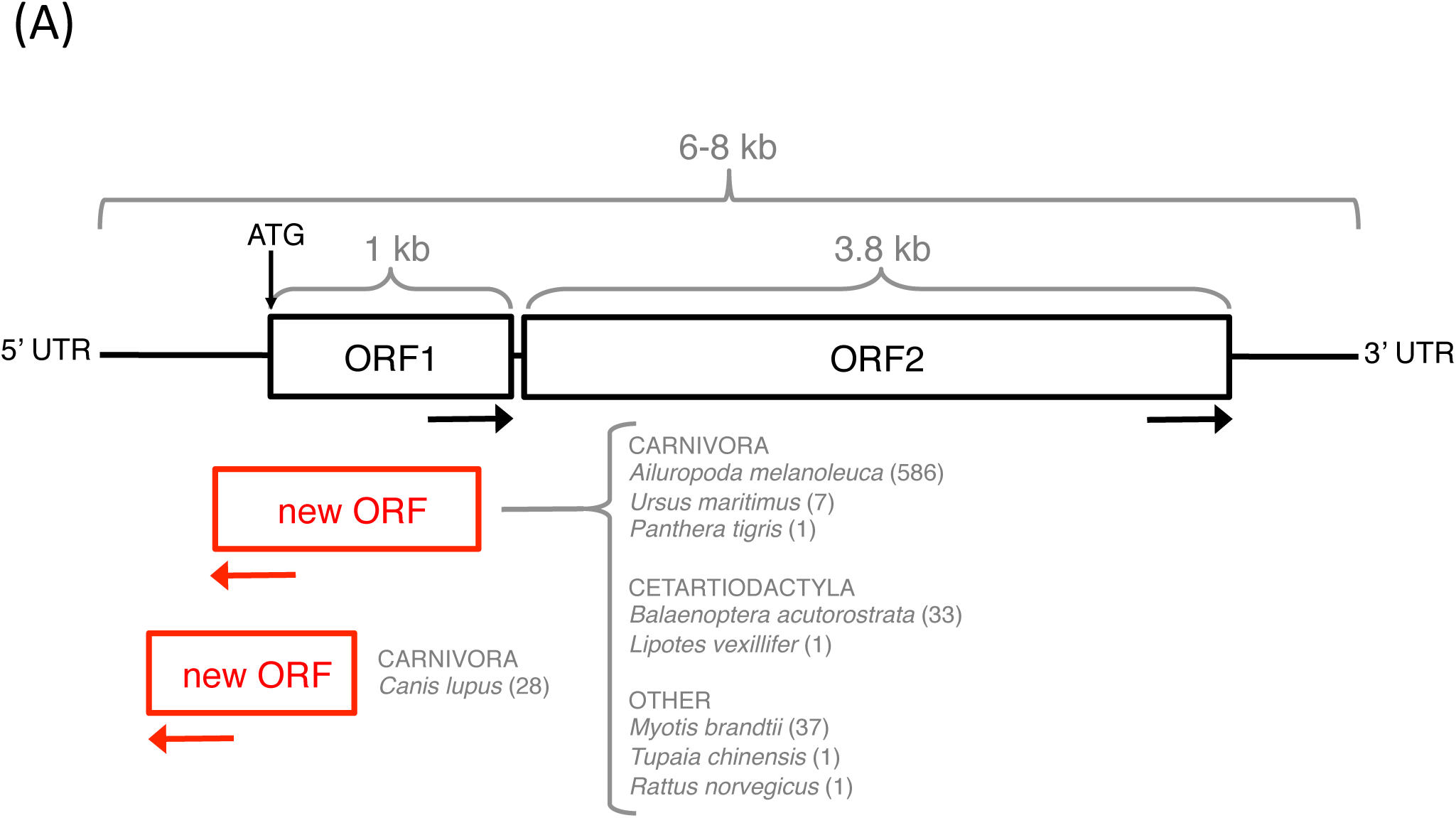
Novel antisense open reading frames found in some mammals. **Fig. 10a. Characteristics and distribution of the antisense ORFs** The position and approximate size of the novel antisense ORFs, as well as the order/species they are found in and the number of L1s that contain this ORF (in brackets). Panda (*Ailuropoda melanoleuca)* has by far the most. The antisense ORFs found in the dog genome (*Canis lupus)* met the criteria of overlapping ORF1 and being at least 800bp in length, but were distinct in that they overlapped only the 5′ end of ORF1. These ORFs have no known functional domains.

Using the same procedure as previously described for ORF1p and ORF2p (Supp. Figure 5), we extracted and aligned the reverse ORF proteins in each species to generate a representative consensus sequence, then aligned the consensus sequences and inferred a maximum likelihood phylogeny (Figure 10b). Apart from the ORF proteins from dog *Canis lupus* (which seems to be a distinct type of reverse ORFp) and Siberian tiger *Panthera tigris* (based on a single sequence rather than a consensus), the other reverse ORF proteins seem to uphold expected species relationships. Scanning the ORF proteins against the current Pfam database using HMMer (Finn et al. 2011) revealed that the 37 *Myotis brandtii* ORFp (only eight of which are unique) identify as Pico_P1A: a picornavirus coat protein (VP4). The dog and minke whale ORFp also got hits with HMMer, but mostly as uncharacterised protein domains.

**Fig. 10b.**
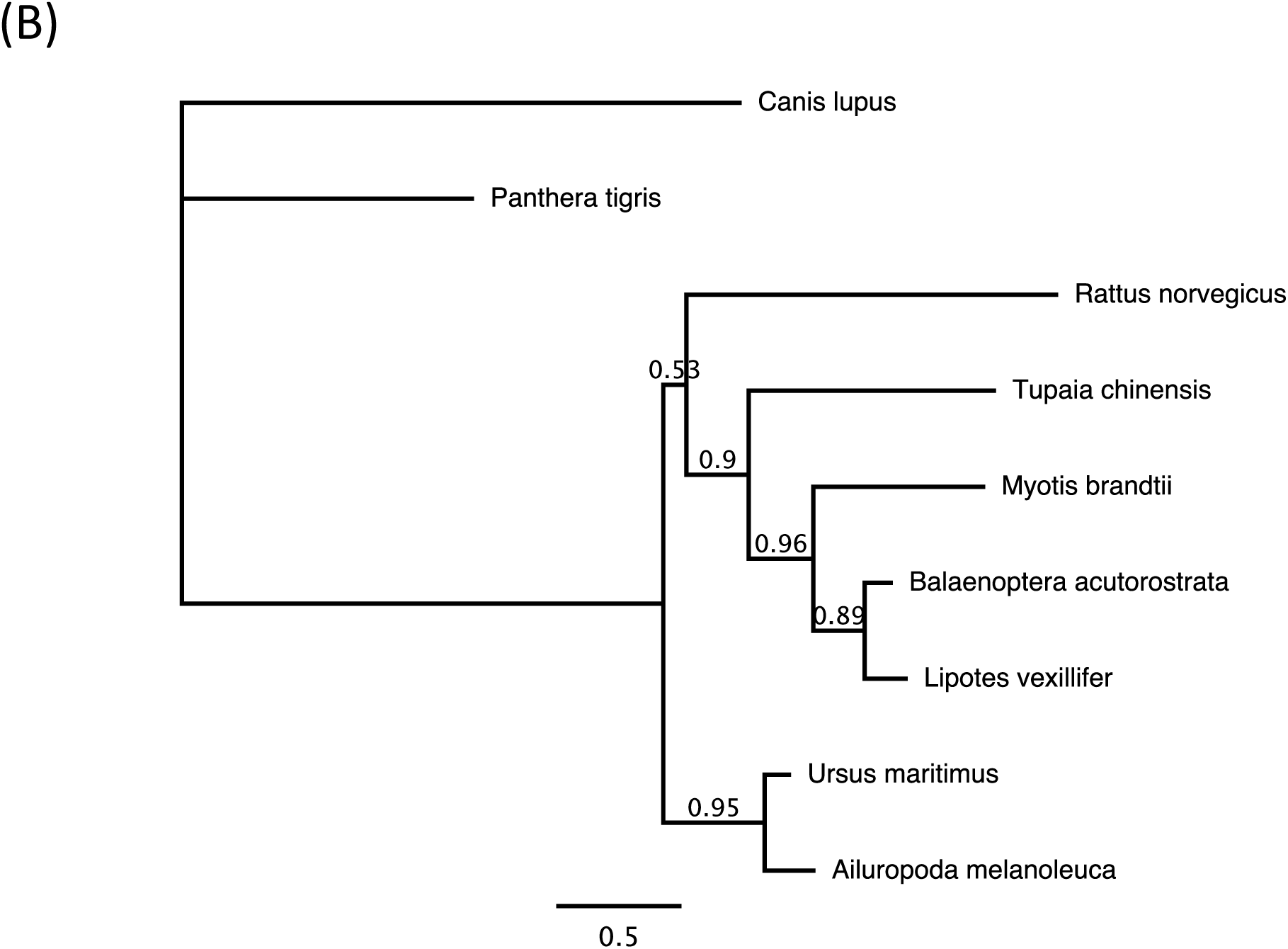
Antisense ORFp species consensus tree. Maximum likelihood phylogeny inferred using FastTree from extracted and aligned L1 reverse ORFp consensus sequences. Expected species relationships seem to be preserved, excluding *Panthera tigris* (which is based on a single sequence rather than a consensus) and *Canis lupus* (which appears to be a distinct type of antisense ORFp).

## Discussion

### Extinction Of L1s In Mammalian Taxa – Known Vs. New Events

An L1 element is called ‘extinct’ if it completely loses its ability to retrotranspose. If there is very low (but still extant) activity in the genome, this has been referred to as ‘quiescence’ rather than extinction (Yang et al. 2014). Figure 3 shows all of the known cases of L1 extinction (not quiescence) out of the 98 mammalian species analysed in this study: three pteropodid bats (Cantrell et al. 2008, Yang et al. 2014) and the thirteen-lined ground squirrel *Ictidomys tridecemlineatus* (Platt and Ray 2012). Other confirmed cases of L1 extinction include the spider monkey (Boissinot et al. 2004) and all studied Sigmondontinae rodents except for the Sigmodontini tribe (Casavant et al. 2000, Grahn et al. 2005), which were not included in this study because there are no public genome assemblies available.

Novel L1 extinction species candidates found in this study include three primates, eight rodents, one lagomorph, seven cetartiodactyls, three carnivores, two perissodactyls, four bats, three Insectivora, five Afrotherian mammals and two marsupials (Figure 3). To our knowledge, these species have not been previously investigated as L1 extinction candidates (although some closely related species have been, e.g. *Peromyscus californicus* (Casavant et al. 1998)).

Evidence of a retro-element extinction event is often difficult to confirm, because we cannot determine whether it occurred in the individual genome or at the species level. The easiest extinction event to observe is one that is ancestral, such that a large monophyletic group of species all lack evidence of recent L1 activity (Grahn et al. 2005). For example, Cantrell et al. (2008) confirmed L1 extinction of the Pteropodidae megabat family by showing that the event had been inherited in 11 sampled genera. Based on this criterion (with each group consisting of at least two different species), there are six potential extinction events shown on our phylogeny (marked ‘**?**’ on Fig. 3): the Delphinidae family in Certartiodactyla; a subgroup of carnivores containing ferret, walrus and Weddell seal; the Rhinolophoidea superfamily of bats – which is closely related to the Pteropodidae family; all three species in the order Eulipotyphla; Afrotherians (except for Proboscidea and Hyracoidea); and metatherians (excluding Didelphimorphia).

Returning to the L1 extinction candidate species which appear paraphyletic or polyphyletic, we see that there are several possible explanations for these occurrences. First, these may be individual organism-specific changes - as with the extinction of L1s in the ground squirrel, which corresponded to a steady decline of all TE classes in that genome (Platt and Ray 2012). Second, the re-emergence or persistence of L1 activity in closely related species suggests that these are examples of quiescence rather than extinction. This may especially be true for the primates, where we already know of an extinct/quiescent species (spider monkey) (Boissinot et al. 2004) and we see that even the active primates have very few active L1 copies (barring obvious exceptions like human and snub-nosed monkey). Such a scenario suggests that there is a fine line between calling an L1 active or extinct, and a lot of these primates may have only recently become inactive. This theory is supported by the observation that many of the primate L1s contain truncated (and thus inactive) ORF2 sequences. Lastly, it is possible that these supposedly extinct species appear so because of the draft quality of the genome assembly used. Experimental analyses such as *in situ* hybridisation would be needed to confirm complete loss or presence of L1 activity (Grahn et al. 2005, Cantrell et al. 2008).

### The Difference between Retrotransposition Potential and Activity

The majority of this study focuses on identifying L1 elements that have retrotransposition potential, and therefore may be active within the genome and causing change. But what does it mean for an L1 to be active? We can label an element as having the ‘potential’ to be active by looking for intact open reading frames, or calculating the proportion of intact full-length L1s in the genome. But to be truly ‘active’, the element must provide evidence that it is doing something in the genome, not just that it has the potential to. So for L1 elements, effective activity should be confirmable by substantial replication and propagation of the element throughout the genome.

The distribution of L1 activity shown in Figure 5 clearly illustrates this concept. There are three things that are immediately obvious in this figure: (1) non-mammalian animal species (shown in red) and plant species (e.g. green alga) have a surprisingly high percentage of active elements but low copy number; (2) the majority of mammals have a huge number of active L1s, but a consistently low (< 20%) active/non-active ratio; (3) several mammalian species (e.g. minke whale, antelope, snub-nosed monkey, mouse, sheep) stand out because they have a high active percentage, unlike the other mammals. The variation between species illustrates those that are potentially active versus those that are truly active.

Addressing the first of these observations – non-mammalian species (plants and animals) all seem to have a relatively low L1 copy number. This is not unexpected in itself; many of these elements are divergent and have accumulated mutations, suggesting that they are older than their mammalian counterparts (as shown by the longer branch lengths in Figure 8). What is surprising is that, based on the identification of intact ORFs, the majority of L1s in these genomes seem putatively active. All ten of the non-mammalian animals have more than 19% of their full-length L1s active. Furthermore, green alga (*Coccomyxa subellipsoidea*) is the only species with 100% of its L1s apparently active. But are these L1s really active? Such low copy number would suggest that there is high retrotransposition potential, but low activity or effectiveness. Screening of the non-mammalian genomes for partial L1 hits using CENSOR (Kohany et al. 2006) revealed very few hits > 2kb, providing further evidence that the L1s in these species are not currently undergoing retrotransposition.

In contrast, we know that mammalian species typically have a high L1 copy number (Lander et al. 2001, Mouse Genome Sequencing et al. 2002). We also know that L1 retrotransposition is thought to be extremely inefficient because the vast majority of new insertions are 5’ truncated and thus inactive (Sassaman et al. 1997, Boissinot et al. 2000). This seems to be the case for most of the mammals analysed in this study: although they have a high number of active L1s, the number of inactive L1s is much greater (~80%); hence they have a low level of observable activity within the genome.

However, there are a few mammals that have both a high L1 copy number and a high active percentage in the genome. Indeed, the most significantly ‘hyperactive’ species (minke whale) has never been mentioned before in the context of L1 activity, yet it contains 3534 active L1s that make up more than 67% of the total full-length L1 content in the genome – far surpassing the retrotranspositional activity of mouse. This directly contradicts the belief that most full-length L1s are inactive or truncated during replication. As such, it is a good indication that these species are truly active, not just potentially active. These L1s are dynamically replicating and expanding within the genome, resulting in a large copy number of elements that share high pairwise identity with each other. Therefore, out of the 125 putatively active species found in this analysis, these five genomes would be the best model organisms for studying genomic change due to L1 retrotransposition.

### The Myth Of The Master Lineage

The master lineage model is an evolutionary scenario where the active elements in a genome give rise to a single active lineage that dominates long-term retrotransposition (Clough et al. 1996). Phylogenetic analyses such as dendrogram constructions are often used to give an indication of existent lineages (Grahn et al. 2005, Adelson et al. 2009), under the rationale that longer branch lengths represent accumulated mutations (including insertions and deletions) due to age, whereas shorter branch lengths signify younger, closely related elements with little nucleotide divergence from the master template. If all of the active elements form polytomies with very short branch lengths, as opposed to multiple divergent clusters, then this would be an example of a strict master lineage model.

It is hypothesised that there is selective pressure for the master LINE (and/or SINE) lineage to monopolise active retrotransposition in mammalian model organisms (Platt and Ray 2012). Our data supports this – all of the ‘hyperactive’ species and many of the potentially active ones contain a single active L1 family/cluster, as shown in Figure 6 with the snub-nosed monkey example. This seems somewhat counterintuitive; given the vast number of active elements, it should be feasible for numerous independent lineages to amplify, over time. A possible explanation is that the single lineage we observe is due to a master element which was particularly effective at evading host suppression mechanisms, and thus initiated widespread retrotransposition throughout the genome.

In some species with relatively low active copy number, such as *Myotis brandtii* (Figure 7), there appear to be multiple simultaneously active lineages. A similar situation was observed in the (now extinct) megabat L1s (Yang et al. 2014) and two putatively active L1 lineages in rodent *Peromyscus californicus* (Casavant et al. 1998). There are various theories as to how multiple lineages may arise: e.g. after a period of low activity, multiple ‘stealth driver’ (Cordaux and Batzer 2009) elements may be driven to retrotranspose at the same time; or horizontal acquisition of a retroelement from a different species can produce a foreign active lineage alongside the native lineage. Nonetheless, not much is known about how both lineages can be maintained, if there really is selective pressure to adhere to a master model. Yang et al. (2014) speculate that if the lineages are specialised on different tissue types (e.g. male germ line vs. female germ line), they can co-exist without competition – however, this is countered by the observation that in mouse, most L1 retrotransposition events seem to occur in the early embryo rather than in germ cells (Kano et al. 2009). Furthermore, the fact that we do not observe any high copy number species harbouring more than one lineage suggests that multiple lineages are inhibitory to retrotransposition: either through competition, or because it increases the chance that both lineages will be detected and suppressed by regulation mechanisms, so neither lineage can effectively proliferate within the genome.

### Discordance between ORF Nomenclature and Domain Classification

A predictable side effect of having access to more data and discovering new domains is that the existing nomenclature may need revision to reflect this new information. Based on the existing Type system for ORF1p elements (Khazina and Weichenrieder 2009), 77 plant species and 6 non-mammalian animals (five mosquito species and sea squirt *Ciona savignyi*) belong to Type I, due to the presence of RRM and/or zf-CCHC domains (Figure 9a,b); and the single remaining plant species (*Coccomyxa subellipsoidea*) belongs to Type V: unclassified ORF proteins. Such a categorisation is potentially misleading since it implies that Type I sequences are alike and share high amino acid similarity – and even the HTH_1 domain in *C. subellipsoidea* cannot be that distantly related, by virtue of it being an ‘ORF1p’. But at what point does a domain variant become too different to be an ORF1p? To demonstrate this, the clustering analysis in Figure 9c was performed using default settings where two proteins in a pair are included in the same family if the homologous segment pairs have at least 50% similarity over 80% coverage (Penel et al. 2009). Even at this low threshold, we see that the ORF1 sequences (all of which supposedly belong to ‘Type I’) form completely independent clusters. Construction of the ORF2p consensus tree (Figure 8) revealed that plant ORF2p domains are also diverse, with very low pairwise identity between sequences (Supp. Table 10).

**9c).**
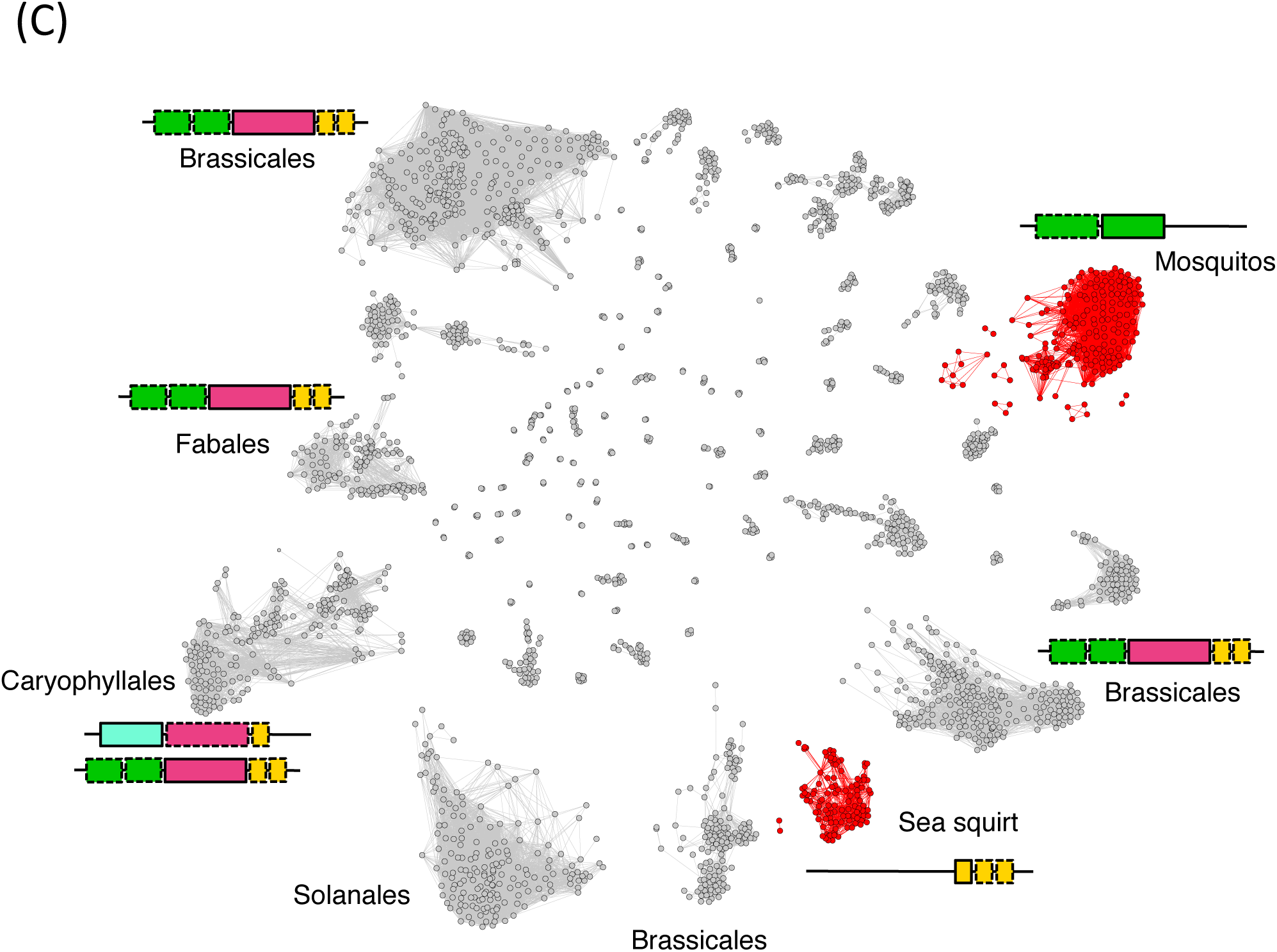
Network clustering analysis of Type I ORF1 proteins. The clustering was performed with default parameters (two proteins are included in a family if the homologous segment pairs share at least 50% similarity over 80% coverage). All of the proteins used are loosely categorised as Type I, yet we see that they form distinct clusters. Even Type I ORF1p within the same order (e.g. Brassicales) do not all cluster together. Non-mammalian animal species are coloured red, plants are coloured grey.

Accordingly, we propose a more informative revision to the nomenclature to refer to ORF proteins by the dominant functional domain: e.g. ORF2p = RVT_1-ORFp for mammals (RVT_3-ORFp for some plants – see Supp. Table 13); ORF1p = HTH_1-ORFp for *C. subellipsoidea*. This allows us to forego predetermined Type or ORF# labels, especially for unusual cases. The discovery of additional ORF proteins such as the primate-specific ORF0 (Denli et al. 2015) or the reverse ORFp found in this study (Figure 10) makes a compelling argument for re-naming.

### Confounding Bias Due To Genome Assembly Quality

Advances in technology mean that genomes are now being sequenced at alarmingly fast rates. However, once sequenced, many genomes tend to remain in their error riddled, scaffolded state. The majority of genomes used in this study are draft assemblies, so it is important to check that the quality of the assembly is not affecting the results (either by restricting the ability to detect repetitive 6kb elements, or by creating false positive hits from misread errors). Accordingly, analysing independently-assembled closely related species (e.g. within the same genus or even species) can act as a quality control. For example, three horse genomes were included in this study: *Equus przewalski* (submitted by IMAU, contig N50 of 57,610, SOAPdenovo assembly method used), *Equus caballus* Thoroughbred (submitted by GAT, contig N50 of 112,381, ARACHNE2.0 assembly method used) and *Equus caballus* Mongolian (submitted by IMAU, contig N50 of 40,738, SOAPdenovo assembly method used) (see Supp Tables 1, 2, 3). Based on the submitter, contig N50 and assembly method, *Equus przewalski* and the Mongolian *Equus caballus* would be expected to be the most similar. Based on species relationships, one would expect the two *Equus caballus* horses to be more similar. However, the actual findings show that while all three horses exhibit L1 presence, only *Equus przewalski* and *Equus caballus* (Thoroughbred) are still potentially active. This is interesting because it suggests that (a) Mongolian horse has undergone an individual L1 extinction event, and (b) using the same submitter and assembly method did not skew the results. As a contrasting example, the three *Arabidopsis* species that were submitted independently (*A. halleri:* TokyoTech, *A. lyrata:* JGI, *A. thaliana:* Arabidopsis Information Resource), have very different contig N50 values (*A. halleri:* 2864, *A. lyrata:* 227,391, *A. thaliana:* 11,194,537) and used different sequencing strategies (*A. halleri:* Illumina, *A. lyrata:* Sanger, *A. thaliana:* BAC physical map then Sanger sequencing of BACs) have very similar results in terms of L1 presence, activity and open reading frame structure. In fact, Illumina seems to be the most widely used sequencing technology across all the genomes (mammalian, non-mammalian and plant) but it does not appear to introduce platform specific artefacts and affect the L1 analysis with respect to the different evolutionary patterns of the different species groups.

The assembly level does not seem to influence the results either: out of the five so-called ‘hyperactive’ mammalian species labelled in Figure 5, three (minke whale, snub-nosed monkey, antelope) are scaffold-level assemblies, while two (mouse and sheep) are chromosome-level with noticeably higher N50 values. One might argue that this just shows that draft assemblies are more likely to have duplication or misread errors, leading to greater L1 copy number. However, an all-inclusive de-duplication of the 338,204 combined L1 sequences showed that 337,732 sequences are unique (only 472 duplicates in the whole 503-genome dataset), and the largest cluster of duplicates had 16 elements (from *Fukomys damarensis*). Moreover, all of the L1 sequences have different flanking regions, suggesting that they are more likely to be true duplicates than assembly errors.

### Implications For Our Perception Of Genome Evolution

This study complements those of Khan et al. (2006) (primates), Sookdeo et al. (2013) (mouse), Yang et al. (2014) (megabats), Metcalfe and Casane (2014) (Jockey non-LTR elements) and Smyshlyaev et al. (2013) (plants) in demonstrating the diversity of TE evolutionary patterns across species. We have identified L1 sequences from 503 different genomes, including 28,967 ORF1 and 17,952 ORF2 proteins with distinct domain variations that strain the current L1 classification system. While most mammals and plants still exhibit some form of L1 activity, L1 extinction/quiescence in modern species appears more widespread than previously believed – with the discovery of new extinction candidates leaving us better equipped to identify common factors in the genomic landscape that contribute to TE suppression. Conversely, investigation into ‘hyperactive’ species such as minke whale and snub-nosed monkey, whose retrotranspositional activity seems to far surpass that of human, rat and mouse, could be used to study the extent to which L1s cause genomic change. Perhaps the presence of reverse ORFs helps the L1 in these species to attain hyperactivity. As always, it is likely that our findings here are only the very tip of the iceberg. We present this data with the hope that it will provide a definitive reference for future studies, aiding our understanding of eukaryotic evolution.

## Acknowledgements

We would like to thank our collaborators (Broad Institute of MIT and Harvard for the chromosome-level elephant genome, and Frank Grutzner from the University of Adelaide for the echidna assembly) for making private genome assemblies available to us. We would also like to thank Iain Searle for taking the time to read the manuscript in full and offer helpful comments. Finally, this paper would not be possible without the invaluable insights of Lu Zeng, R language mastery of Reuben Buckley and HMMer knowledge of Zhipeng Qu.

